# Mechanoregulation of biofilm architecture promotes *P. aeruginosa* antibiotic tolerance

**DOI:** 10.1101/2022.02.16.480709

**Authors:** Alice Cont, Joseph Vermeil, Alexandre Persat

## Abstract

In the wild, bacteria are most frequently found in the form of multicellular structures called biofilms^1^. Biofilms grow at the surface of abiotic and living materials with wide-ranging mechanical properties. Despite their co-occurrence during infection, we still lack a clear understanding of how mechanics regulate biofilm architecture and the physiology of resident bacteria. The opportunistic pathogen *Pseudomonas aeruginosa* forms biofilms on indwelling medical device^2^ and on soft tissues including burn wounds and the airway mucosa^3^. Here, we demonstrate that mechanical properties of hydrogel material substrates define *P. aeruginosa* biofilm architecture. We show that hydrogel mesh size regulates twitching motility, a surface exploration mechanism priming biofilms, ultimately controlling the arrangement of single cells in the multicellular community. The resulting architectural transitions increase *P. aeruginosa*’s tolerance to colistin, a last resort antibiotic. Our results thereby establish material properties as a regulator of biofilm architecture and antibiotic efficacy.

---

Bacteria assemble into biofilms by attaching to surfaces and subsequently dividing while embedding themselves in a self-secreted matrix^4,5^. In this process, biofilms generate internal forces that deform soft substrates, which in turn can disrupt host epithelial tissue^6^. On the other hand, and as one might suspect, mechanical forces play a role in bacterial attachment, surface-specific motility and biofilm organization^7,8^. Material properties such as topography, chemistry, surface charge, hydrophobicity are known physicochemical mediators of biofilm formation, mainly impacting initial bacterial adhesion^9–14^. Explorations of bacterial adhesion on elastomers and hydrogels suggest that material mechanical properties may mediate biofilm formation^15–17^. Even though these studies show contrasting effects, there is parallel evidence that bacteria mechanically sense and respond to solid surfaces^8^. For example, *P. aeruginosa* mechanosense surfaces to regulate virulence gene expression and surface-specific twitching motility, both of which are important regulators of biofilm formation^18–21^. Due to the prevalence of surface mechanics and mechanosensing in *P. aeruginosa* surface adaptation, we hypothesize that the material properties of material substrates regulate *P. aeruginosa*’s ability to form biofilms.

To investigate the link between substrate mechanics and *P. aeruginosa* biofilm formation, we synthesized thin films of poly(ethylene glycol) diacrylate (PEGDA) hydrogels as biofilm growth substrates. These synthetic polymeric networks are optically clear, biocompatible, have homogeneous mechanical properties and are relatively soft thereby matching properties of living tissues^22^. In addition, the elastic moduli of PEGDA hydrogels can be finely tuned by controlling cross-linking^23^. To investigate the contributions of material properties on biofilm formation, we screened an assortment of PEGDA hydrogels cross-linked from prepolymer of different chain length (700 Da to 6000 Da) and concentrations (10% w/v to 30% w/w). We generated thin hydrogels films (~50 μm) at the bottom surface of microfluidic channels, and subsequently initiated biofilm growth under flow (Figure 1A). *P. aeruginosa* successfully colonized the surface of all hydrogels. We however noticed dramatic difference in the colonization patterns across conditions (Figure 1B). On hydrogel with relatively long and dilute PEG, *P. aeruginosa* predominantly formed well-defined isolated biofilms (Figure 1Bi). These biofilms grew into tall structures with high aspect ratio. In contrast, *P. aeruginosa* populations appeared to colonize the surface of hydrogels from lower molecular weight or higher PEGDA concentrations more uniformly (Figure 1Bii-iv). These biofilms only slightly extended in the depth of the channel, normal to the surface. In the most extreme case of a 20% PEGDA at 700 Da, it became difficult to distinguish biofilm colonies as bacteria rather formed monolayers. In other words, decreasing the polymer chain length and increasing concentration promotes the transition from compact dome-shaped structures with near unity aspect ratio, to flat and spread out biofilms with relatively lower aspect ratio (Figure 1C).

**Figure 1.**
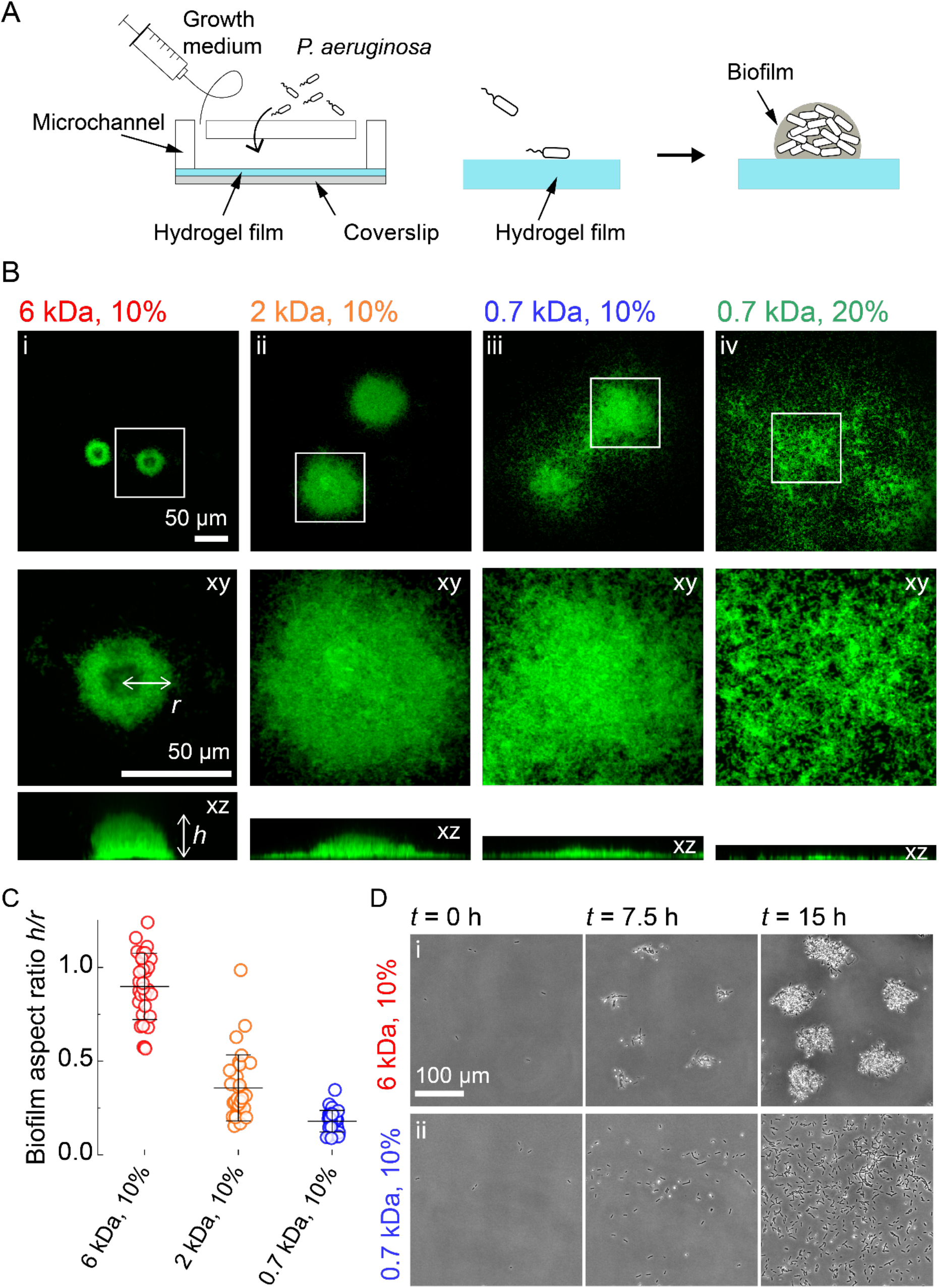
Hydrogel elastic substrates regulate *P. aeruginosa* biofilm architecture. (**A**) Illustration of experimental setup where we generate thin PEG hydrogel films at the bottom surface of microchannels. These devices allow us to study biofilm formation on soft materials at high resolution. (**B**) In-plane and cross-sectional confocal visualizations show different architectures of *P. aeruginosa* biofilms grown on hydrogels with different chain length (molecular weight, MW) and concentration of PEGDA precursors. For (i) MW = 6000 Da,10% wt/vol, (ii) MW = 2000 Da, 10% wt/vol, (iii) MW = 700 Da, 10% wt/vol, (iv) MW = 700 Da, 20% wt. (**C**) Quantification of biofilm height (*h*) and radius (*r*) on different hydrogels shows an increase in aspect ratio *h/r* with increasing chain length. Each circle corresponds to one colony, black bars represent mean and standard deviation across all values. (**D**) Timelapse visualization reveals that hydrogels regulate *P. aeruginosa* biofilm architecture by regulating initial steps of biogenesis. (i) Biofilms form by division and local attachment in the vicinity of founder cells on long chain PEGDA hydrogels. (ii) Single cells explore the surface of short chain PEGDA hydrogels to form biofilms by aggregation.

Given prior observations, we anticipated that substrate stiffness could regulate biofilm biogenesis^17^. We were however surprised that biofilms adopted strikingly different architectures on PEGDA hydrogel with nearly identical moduli (Figure 1B, Table S1). We therefore investigated the mechanisms by which material properties regulate biofilm architecture. Timelapse visualizations revealed noticeable differences between materials at the early stages of biofilm formation. Microcolonies rapidly appeared in the first few hours of growth on PEGDA hydrogels with high molecular weight (Figure 1Di, movie 1). On shorter chain PEGDA, we observed most *P. aeruginosa* cells exploring the surface as they grew and divided, leading to a more uniform distribution on the surface (Figure 1Dii, movie 1). While this mechanism tends to decrease local cell density near founder cells, it still promotes bacterial aggregation on longer timescale, allowing the development of multicellular structures. These visualizations suggest that distinct hydrogels control biofilm architecture by regulating initial surface exploration.

To nucleate biofilms, *P. aeruginosa* first attaches to and navigates on surfaces using twitching motility, which ultimately drives cellular aggregation^24^. Long and thin retractile protein filaments called type IV pili (T4P) power twitching motility. T4P propel single cells forward by successively extending, attaching to the surface and retracting. How substrate mechanical properties regulate twitching motility remains unclear, but theory predicts cells pull themselves more efficiently on stiffer materials^25^. Guided by our observations of early biofilm formation, we hypothesized that material properties regulate twitching motility, thereby leading to the different biofilm architectures. To test this hypothesis, we performed extensive measurements of *P. aeruginosa* twitching at the single cell level on PEGDA hydrogels. We recorded the trajectories of hundreds of cells per condition, and computed mean speed for each cell along their track (Figure 2 A-C, movie 2). Single *P. aeruginosa* cells barely migrated on PEGDA hydrogels that favored the formation of defined dome-like biofilms (Figure 2 Ai and C). In contrast, cells were much more motile on hydrogels that favored the formation of biofilm monolayers (Figure 2 Aii and C). Thus, the final architecture of a biofilm reflects the motility patterns observed on the distinct hydrogels. On substrates inhibiting motility, *P. aeruginosa* cells divide and accumulate near the initial founder cells, forming tightly packed dome-shaped biofilms. On substrates promoting motility, cells disperse on the surface as they divide, thereby limiting accumulation near founder cells but promoting surface occupation.

**Figure 2.**
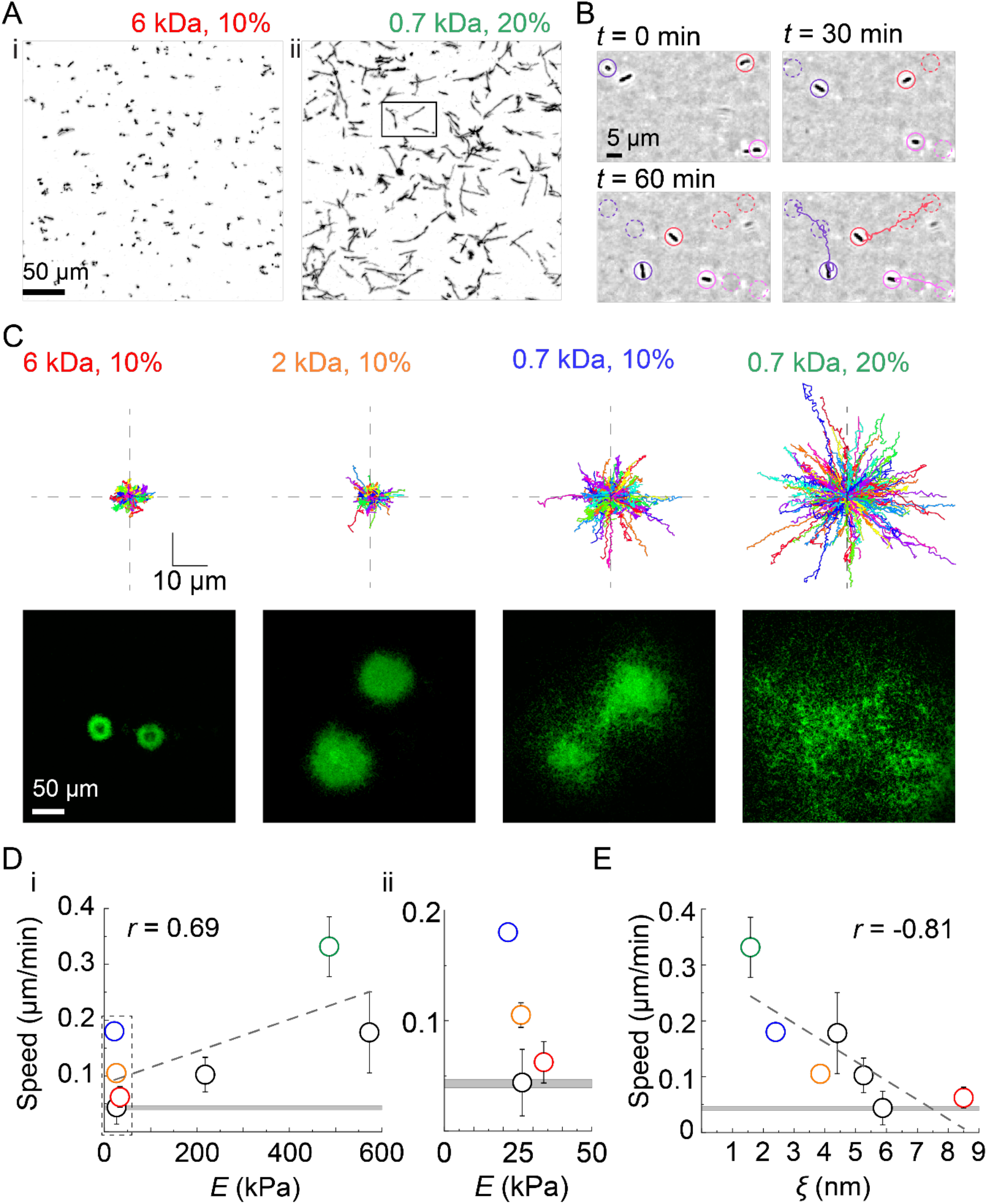
Hydrogel mesh size modulates biofilm architecture by regulating twitching motility. (**A**) Cumulative surface coverage of bacterial trajectories during the first hour of contact with the indicated PEGDA gel. Black signal corresponds to bacterial, white corresponds to unexplored surface. (**B**) Timelapse visualization of *P. aeruginosa* twitching on the surface of PEGDA 700 Da at 20%, and respective trajectories (from selection in Figure 2A-ii). (**C**) Trajectories of 300 randomly selected cells on four different hydrogels and resulting biofilm morphology after 40 h of growth. All trajectories start from the center of the graph. (**D**) Hydrogel mechanical property regulating twitching speed. i) Population mean twitching speed increases with Young’s modulus, but with limited correlation (Pearson correlation coefficient *r* = 0.69). ii) Closeup of twitching speed as a function of modulus at low stiffness. Despite modulus being nearly identical on the four gels, we found large differences in twitching speeds. (**E**) Twitching speed decreases with increasing hydrogel mesh size, with relatively stronger correlation (Pearson correlation coefficient *r* = −0.81). For (**D**, **E**) dashed lines indicate the linear fit of the data. Circles represent the mean across biological replicates and black bars represent standard deviation (SD). The horizontal grey line represents the mean speed across biological replicates for a T4P retraction-deficient mutant (*ΔpilTU*), and the thickness of the line corresponds to the SD.

How does the material substrate control twitching motility? Consistent with our initial qualitative observations of biofilm architecture (Figure 1B), we only found a slight correlation between PEGDA Young’s modulus and twitching speed (Figure 2Di). At equal polymer chain length, increasing stiffness with larger precursor concentration sped up twitching. However, this trend failed across gels with same concentration but different chain lengths: we measured a 3-fold change in twitching speed on hydrogels of nearly identical moduli (Figure 2Dii). Given the importance of cell and T4P attachment in twitching motility, we reasoned that hydrogel surface topology may affect *P. aeruginosa*’s motility. To estimate substrate density at the surface, we measured the mesh size of the hydrogel films using equilibrium swelling theory. Twitching speeds showed a stronger, negative correlation with mesh size across concentrations and chain lengths (Figure 2E). This revealed that hydrogel mesh size controls twitching motility of single *P. aeruginosa* cells. The strength of adhesion of bacterial cell bodies is indistinguishable between these different mesh sizes (Supplementary figure 1). We thus propose a model where the likelihood of T4P attachment depends on mesh size, thereby controlling the rate of productive T4P retractions. Consistent with this, twitching speeds increase on hydrogels with mesh sizes below 5 nm, a dimension that corresponds to the diameter of a T4P fiber^26^. Overall, our results suggest that T4P attachment to the hydrogel substrate with larger mesh sizes is less frequent, limiting the efficiency of force transmission during retraction.

The biofilm lifestyle is a major contributor of human chronic infections due to its resilience against antibiotic treatments^27–29^. Chronically infected patients are subject to lifelong *P. aeruginosa* infection even under strong antibiotic therapy, which favours the emergence of antibiotic resistant mutants. Multiple bacterial physiological factors improve tolerance to antibiotics. In biofilms, matrix impermeability, metabolic state of residents and increased cell density all contribute to protecting single bacteria from antibiotic stress^27–29^. We however know very little about how environmental factors influence the sensitivity of *P. aeruginosa* to antibiotics by regulating biofilm formation. In light of the distinct biofilm architectures observed on PEGDA hydrogels, we hypothesized that antibiotic efficacy could differ as a result of material properties. We thus tested the efficacy of colistin, a last resort antibiotic against *P. aeruginosa* infections. We grew biofilms on the different hydrogels for 46 h and subsequently challenged them with colistin for 1 h. To test antibiotic efficacy, we measured the volume of live biomass after treatment using a live-dead fluorescent stain. After 1h of colistin treatment, 50% of the population was killed for biofilms growing on smaller mesh size hydrogels (Figure 3A). This proportion was reduced to 40% on intermediate hydrogels, and went as low as 20% on the lowest mesh-size hydrogels, highlighting a strong decrease in drug efficacy. High resolution confocal images revealed distinct spatial patterns of colistin killing (Figure 4B). On small pore size gels, bacteria were killed uniformly, irrespective of their position in the biofilm. In contrast, biofilms grown on larger mesh size hydrogels showed heterogeneity in cell death. Bacteria at the outer edge of colonies (rim, R) were mostly dead, while cells at the core (C) of biofilms remained largely unaffected by colistin treatment (Figure 4C). Given the spatial pattern of killing, we reasoned that such differences were due to colistin transport into the biofilm. We thus computed the surface to volume ratio in each architecture. Defined biofilms growing on more porous gels have lower surface to volume ratio than less porous gels (Figure 4D). Thus, biofilms growing on more porous gels are more tolerant due to longer diffusion times towards the biofilm core. Treatment on the less porous gels is more efficient as single cells are further exposed to the surrounding fluid.

**Figure 3.**
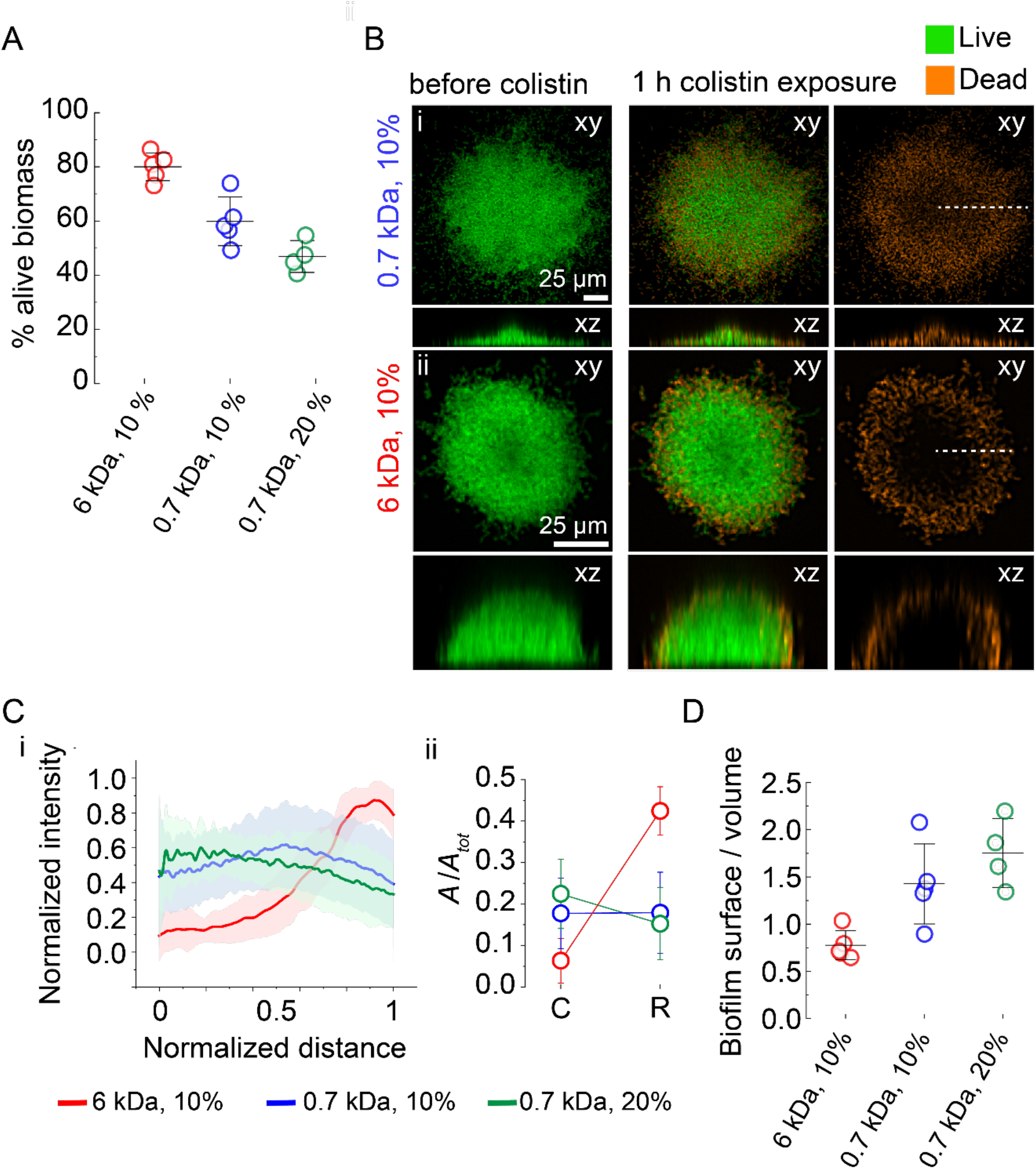
Mechanoregulation of biofilm architecture promotes *P. aeruginosa*’s tolerance to antibiotics. (**A**) Quantification of live biomass after a 1 h colistin treatment, relative to the total biomass before antibiotic exposure. Biofilms grown on hydrogels with larger mesh size are more tolerant to colistin. (**B**) In-plane and cross-sectional confocal visualizations shows differences in colistin killing patterns. (**C**) (i) Dead stain fluorescence intensity profiles computed along a biofilm radius (dashed lines in Figure 3B) highlight a more uniform distribution of dead cells in biofilms grown on hydrogels with smaller mesh size. Distance is normalized to each biofilm radius. Lines represent the average across single colonies, and shaded area correspond to the standard deviation. ii) Integrated normalized area under the curves in the biofilm core (C, between 0 and 0.2 distance unit) and the biofilm rim (R, between 0.8 and 1 distance unit). Circles represent the integrated normalized area and error bars the SD from the curve in shown in panel i. (**D**) Biofilm surface-to-volume ratios grown on different gels. Lower surface to volume ratio decreases overall exposure of single cells to antibiotics, which represses bacterial killing and increases tolerance to colistin. For (**A**, **D**) Circles represent the mean value for each chip, black bars represent mean and error bars the SD across these values.

**Figure 4.**
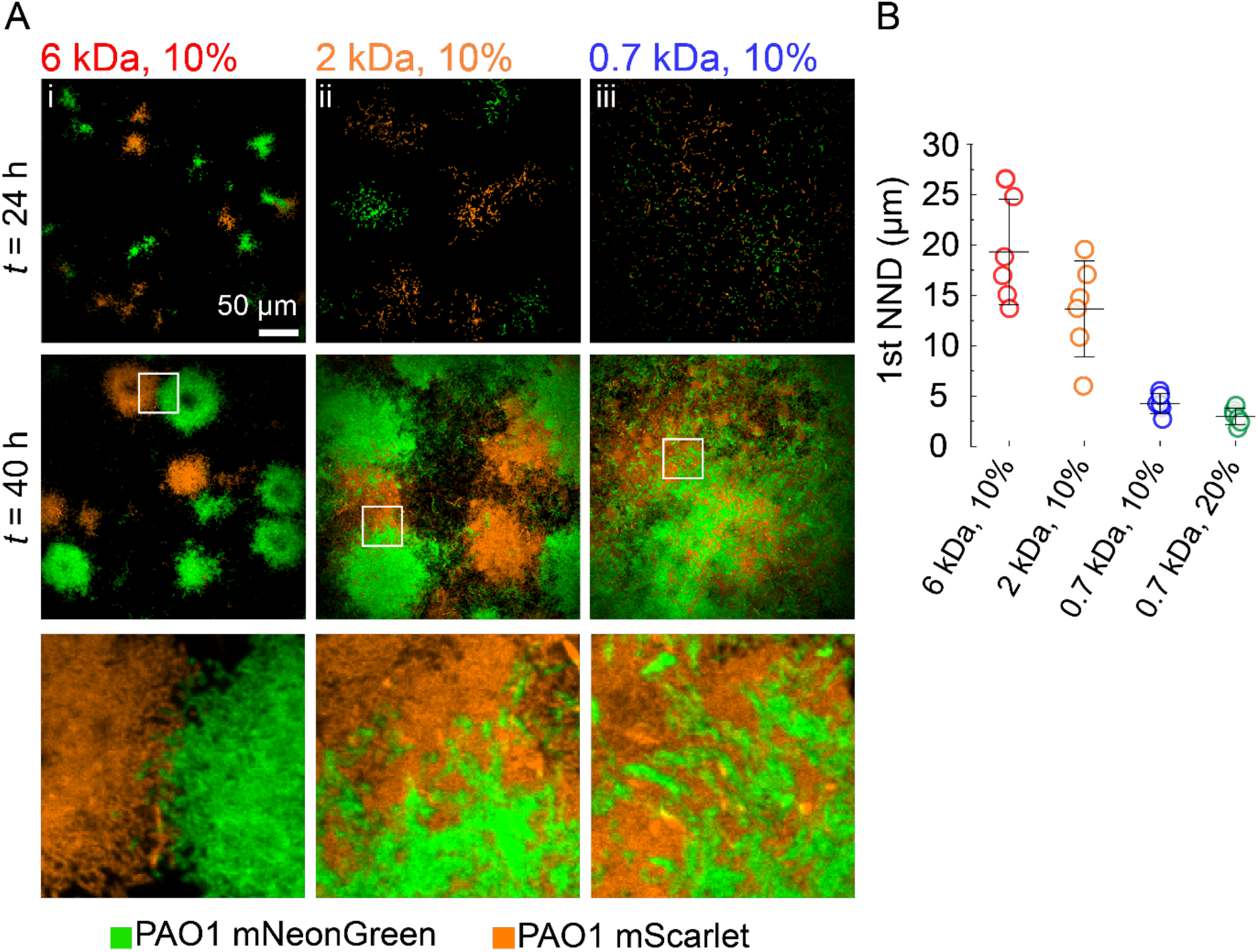
Hydrogel substrates regulate the spatial organization of heterogeneous *P. aeruginosa* biofilms. We grew biofilms from a mixture of *P. aeruginosa* clones constitutively expressing mScarlet (orange) or mNeonGreen (green) (**A**) In-plane confocal visualization at 24 h and 40 h of heterogeneous biofilms on hydrogels with identical modulus but different mesh sizes. Larger mesh size promotes the growth of segregated biofilm clusters. In contrast, hydrogels with smaller mesh sizes promotes clonal mixing due to increased surface exploration. The bottom row shows close up views from the regions indicated by white frames. (**B**) Quantification of the mean first nearest neighbor distance (1^st^ NND) between the mNeonGreen and mScarlet clones. Bacteria forming biofilms on small mesh size hydrogels are more prone to encounters with different clones, forcing them to compete. Circles represent the mean value for each chip, black bars represent their mean and error bars their SD.

Single cell movements modulate the spatial organization of heterogeneous biofilms, ultimately governing how different clones or species compete or cooperate^30,31^. In flow, swimming motility tends to disperse *Caulobacter crescentus* biofilm lineages by spreading out the progeny of founder cells ^32^. This in turn affects the mixing of different clones coming from distinct founder cells. By analogy, we reasoned that twitching patterns observed on the different materials may affect the clonal organization by affecting lineage structure. We therefore explored the relationship between hydrogel mechanical properties and the mixing of heterogeneous biofilms. We grew biofilms from mixtures of two wild-type *P. aeruginosa* strains that each constitutively expressed the fluorescent proteins mScarlet and mNeonGreen (Figure 4Ai). On large mesh size hydrogels that inhibited motility, biofilms formed into separate, isolated clusters. Finding *P. aeruginosa* cells of one color within a biofilm of the other was rare. mScarlet- and mNeonGreen-expressing cells were only found in close proximity when biofilms of distinct clones grew sufficiently to touch each other. Clonal lineages became however less segregated as hydrogel mesh size decreased, permitting twitching-dependent dispersion. On intermediate mesh size gels, while biofilms grew into defined colonies, there was a clear mixing between clones (Figure 4Aii). Finally, on the hydrogels with largest mesh size, clones were well mixed (Figure 4Aiii). For each hydrogel condition, we computed the mean first nearest neighbor distance between the mNeonGreen and mScarlet clones (Figure 4B). This distance decreased from 20 μm on large mesh size hydrogels (a length-scale corresponding to the typical radius of a biofilm) to 3 μm on the lowest mesh size hydrogel (corresponding to the size of a *P. aeruginosa* cell). Altogether, we showed that substrate material can have a strong influence on the distribution of genetically distinct bacterial population on the surface. Such material-dependent mixing is of key importance in the fitness of each clone and the evolution of traits that mediate interactions.

Genetic and metabolic factors are known regulators of biofilm biogenesis and architecture^33,34^. Altogether, the cellular mechanisms promoting *P. aeruginosa* biofilms are becoming clearer. However, how these communities form in realistic contexts remains unresolved^3^. In particular, how mechanical signals regulate biofilm formation have been vastly underexplored^8^. Here, by investigating the regulation of biofilm architecture at the surface of defined and tunable hydrogels, we discovered that material properties regulate biofilm architecture. Mesh size, but unexpectedly not stiffness, influence the foundations of nascent biofilm structures by regulating T4P-dependent twitching motility of single *P. aeruginosa* cells. Thus, cells growing on larger mesh size materials have reduced motility, producing tall and dense biofilms that increase population tolerance to antibiotics. This in turn also favors the rise of antibiotic resistant mutants on a longer timescale^27,35^. *P. aeruginosa* may encounter materials of distinct or heterogeneous mesh sizes as it colonizes extracellular matrices at burn wounds or mucus with altered viscoelasticity in the lungs of cystic fibrosis patients. By extension, mechanics could regulate the architecture and antibiotic tolerance of biofilms of other piliated species through a similar mechanism. Looking further, mechanisms involving fimbrae, flagella or other protein receptors in the initial steps of biofilm formation may be affected by material density. Finally, the impact of material properties on the spatial organization of heterogeneous biofilms shows that mechanics could also play a role in social interactions between different species, influencing the relative fitness of bacteria of distinct backgrounds on an evolutionary timescale. Further efforts to understand how biofilms form in realistic physical contexts will reveal the relative contributions of mechanics in biofilm biogenesis, and how they ultimately play a role in infections.

## Acknowledgements

We would like to thank Rok Simic and Nicholas Spencer for performing nanoindentation measurements and for helpful discussions. We are grateful for the financial support provided by the by the Swiss National Science Foundation through the Projects grant number 310030_189084 and the Human Frontier Science Program grant number RGY0077/2020.

---

**Figure S1.**
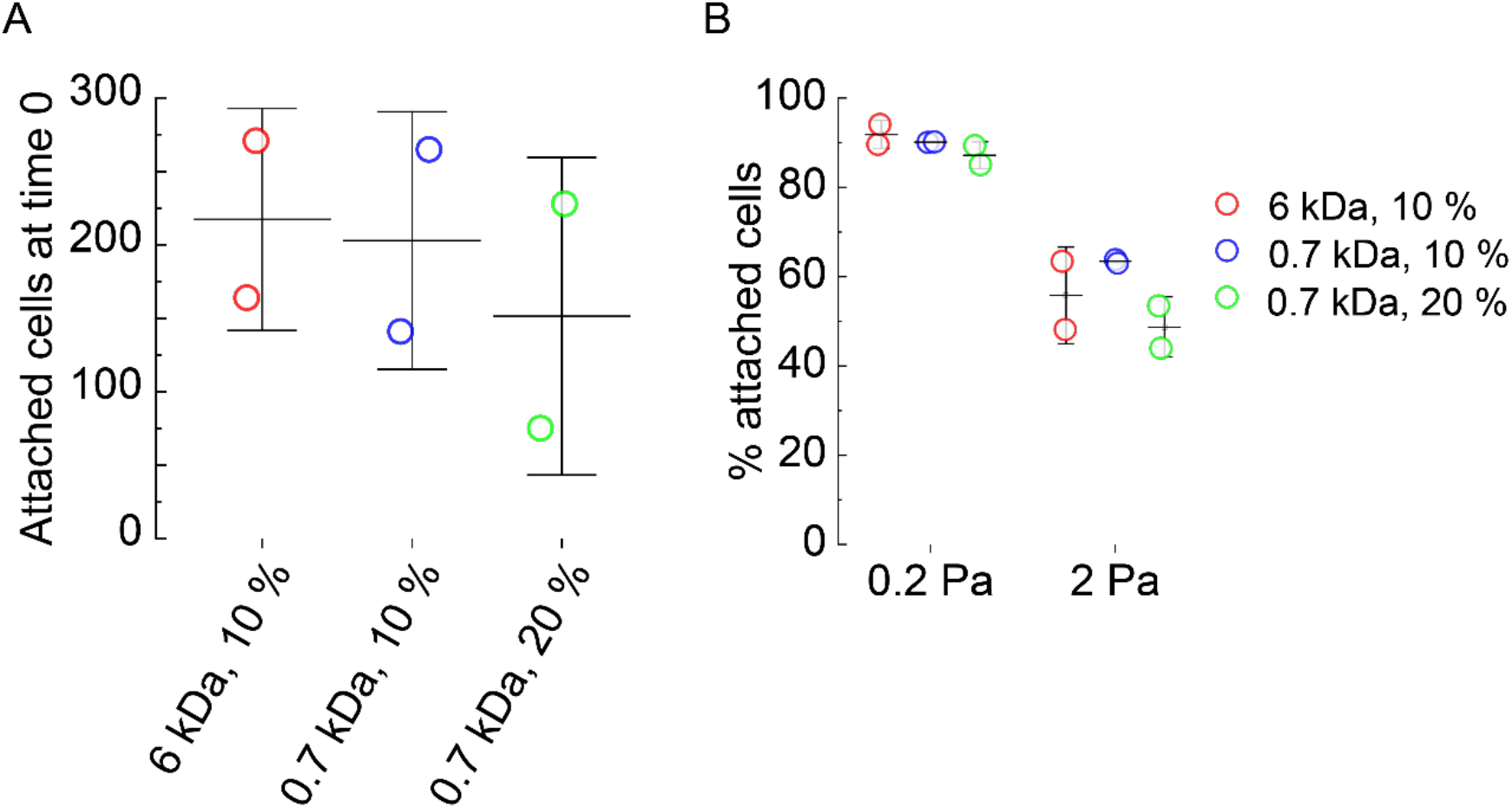
The adhesive strength of *P. aeruginosa* cell body is independent of hydrogel mesh size. (**A**) Quantification of the number of attached *P. aeruginosa* pilus-deficient mutant *ΔpilA* on hydrogels after applying a shear stress of 0.02 Pa. There is no significant difference in bacterial attachment between gel types. (**B**) Percentage of cells that remains attached after applying a shear stress of 0.2 or 2 Pa to cells attached at time 0. Cells are uniformly removed by the flow, independently of hydrogel mesh size. Circles represent biological replicates and black bars represent mean and SD of displayed values.

**MOVIE S1**

Timelapse visualization of *P. aeruginosa* biofilm formation on PEGDA hydrogels with similar modulus (6 kDa, 10% w/v on the left, 0.7 kDa, 10% w/v on the right). Time is in h:min.

**MOVIE S2**

Timelapse visualization of *P. aeruginosa* cells twitching on PEGDA hydrogels (6 kDa, 10% w/v on the left, 0.7 kDa, 20% w/w on the right).

## Methods

### PEG HYDROGELS FABRICATION

#### Solutions

To generate PEG hydrogels we prepared solutions of poly(ethylene glycol) diacrylate (PEGDA) as the precursor and lithium phenyl-2,4,6-trimethylbenzoylphosphinate (LAP, Tokio Chemical Industries) as the photoinitiator in M9 minimal medium (no calcium or magnesium). Different molecular weights (6000 Da, 3400 Da, 2000 Da, 700 Da) and concentrations (10 wt/vol%, 20 wt%, 30 wt%) of PEGDA were used for the formation of the hydrogels (see **Table S1** for details), while the concentration of LAP was kept constant at 2 mM.

#### Hydrogel preparation for mechanical characterization

To measure bulk moduli and mesh sizes of PEGDA hydrogels, we prepared samples by filling cylindrical molds made of PDMS (5 mm diameter, 4 mm height) with the precursor solutions. The molds were covered with a coverslip and hydrogels were formed by irradiating the samples in a UV transilluminator (Bio-Rad Universal Hood II) for 5 min.

#### Hydrogel thin film preparation in microfluidic channels

To obtain thin and flat hydrogel layers, a prepolymer solution was sandwiched between two coverslips. One of the coverslip (25 × 60 mm Menzel Gläser) was cleaned with isopropanol and MilliQ water. The other coverslip (22 × 40 mm Marienfeld) was functionalized with 3- (Trimethoxysilyl)propyl methacrylate (Sigma-Aldrich) to covalently link the hydrogel to the coverslip: we immersed the coverslips for 10 minutes in a solution composed of 1 mL of 3- (Trimethoxysilyl)propyl methacrylate and 6 ml of diluted acetic acid (1:10 glacial acetic acid:water) in 200 mL of ethanol solution 70%. These were subsequently rinsed in ethanol and dried. We then deposited a 30 μL droplet of prepolymer solution on the first coverslip and sandwiched it with the second. The assembly was then placed under the UV transilluminator for 5 min to allow cross-linking. Right after polymerization, the coverslips were separated using a scalpel thereby exposing the hydrogel film surface.

### PEG HYDROGELS CHARACTERIZATION

#### Bulk modulus

The hydrogel cylinders resulting from the polymerization in the PDMS molds were immersed in M9 overnight and tested with a rheometer (TA instruments) in compression mode, at a deformation rate of 20 μm/s. Beforehand, the diameter of the cylinders was measured with a digital caliper, while the height of the cylinder was defined as the gap distance at which the force increases. The elastic modulus corresponds to the slope of the linear fit of the stress-strain curves in the range of 15% strain. The final Young’s modulus is the average modulus of three replicates.

#### Nanoindentation

The hydrogel coated coverslips were immersed in M9 medium right after polymerization. Nanoindentation experiments were performed using an atomic force microscope (AFM, MFP-3D™, Asylum Research, Santa Barbara, USA). The indentation probe was prepared by attaching a silica microsphere (Kromasil, Nouryon - Separation Products, Bohus, Sweden) with a radius of *R* = 11 μm to the end of a tipless cantilever (NSC-36, Mikromash, Bulgaria) with the help of a 2-component epoxy glue (UHU GmbH, Germany). The effective spring constant was calculated as *k = k_0_(L_0_/L)^3^* = 10.01 N/m, where *k_0_* is the spring constant of the bare cantilever, and *L_0_* and *L* are the distances from the base of the cantilever to its tip and to the microsphere, respectively ^1^. The spring constant of the bare cantilever *k_0_* was determined according to the Sader method before attaching the microsphere ^2^. After installing the probe, the laser path was adjusted to center the laser beam on the photodiode and maximize the intensity. The system was then calibrated by pressing the probe against a silicon wafer in water. The force was determined as *F = k x*, and the indentation depth was thus calculated as *d = Z - x*, where *Z* is the vertical piezo displacement and x is the cantilever displacement. The contact with the gel in liquid was determined at the point where the force signal began to deviate more than 2σ from the zero-force line, with *σ* being the standard deviation of the signal noise (~ 20-30 pN). The approach and retraction speeds were set to 1 μm/s. The measurements were performed at 25 °C ± 1 °C. Forty force curves were obtained at different locations of a sample. Elastic moduli were extracted by fitting the Hertzian model to the indentation parts of the measured curves, which showed no adhesion upon the approach ^3^. For the curves that showed a snap-in during the approach, the JKR model was used ^4^.

#### Mesh size

Mesh size was estimated from the equilibrium swelling theory using protocols previously described ^5–9^. For each hydrogel cylinder we determined the volume and the mass in the relaxed (r), swollen (s) and dry (d) states. We measured the volume of the cylinder right after polymerization (*V_r_*) and after immersion in M9 for 24 h (*V_s_*) with a caliper. We then washed the swollen hydrogels in deionized water to remove salts and we dried them overnight in the oven at 80°C. We then measured the mass of the dry network (*M_d_*) and calculated *V_d_* as *M_d_/ρ_PEG_*, with *ρ_PEG_* taken to be 1.18 g/mL. We then calculated the average molecular weight between cross-links, *M_c_* using eq. 1:

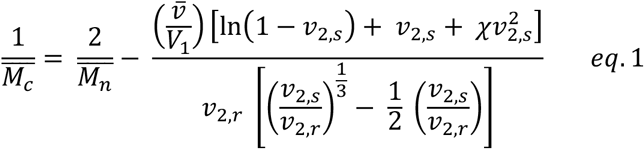

where 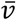 is the specific volume of the polymer (taken to be 0.93 mL/g for PEG), *V*_1_ is the molar volume of water (18 mL/mol), *χ* is the polymer–solvent interaction parameter (taken to be 0.426 for PEG in water), 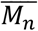 is the average molecular weight of the polymer before cross-linking and *v*_2,*r*_ and *v*_2,*s*_ are the polymer volume fractions:

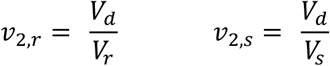

We finally obtained the mesh size ξ with eq.2:

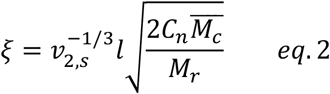

Where *l* is the bond length along the polymer backbone (0.15 nm), *C_n_* is the Flory characteristic ratio (4 for PEG) and *M_r_* is the molecular weight of the repeat unit (44 g/mol).

**Table S1.** Summary of mechanical properties of PEGDA hydrogels.

| PEGDA precursor | Concentration<br>(% w/v or %<br>w/w) | Young's<br>modulus (bulk)<br>± SD (kPa) | Young's<br>modulus (AFM)<br>± SD (kPa) | Mesh size ±<br>SD (nm) |
| --- | --- | --- | --- | --- |
| PEGDA MW 6000<br>(Biochempeg) | 10% w/v | 38.0 ± 13.9 | 33.7 ± 0.8 | 8.50 ± 1.20 |
| PEGDA MW 6000<br>(Biochempeg) | 20% w/w | 265.0 ± 0.3 | 218 ± 6 | 5.26 ± 0.25 |
| PEGDA MW 6000<br>(Biochempeg) | 30% w/w | 470 ± 28 | 573 ± 27 | 4.40 ± 0.14 |
| PEGDA MW 3400<br>(Biochempeg) | 10% w/v | 26.9 ± 1.1 | 26.4 ± 0.6 | 5.88 ± 0.06 |
| PEGDA MW 2000<br>(Biochempeg) | 10% w/v | 37.6 ± 2.1 | 26 ± 0.6 | 3.85 ± 0.17 |
| PEGDA MW 700<br>(Sigma-Aldrich) | 10% w/v | 48.0 ± 1.3 | 21.6 ± 0.6 | 2.40 ± 0.05 |
| PEGDA MW 700<br>(Sigma-Aldrich) | 20% w/w | 750 ± 57 | 486 ± 14 | 1.59 ± 0.03 |

### ASSEMBLY OF HYDROGEL-COATED COVERSLIPS WITH MICROFLUIDIC CHIPS

We fabricated microfluidic chips following standard soft lithography techniques. For biofilm experiments we designed 2 cm-long, 2 mm-wide channels in Autodesk AutoCAD and printed them on a soft plastic photomask. We then coated silicon wafers with the photoresist (SU8 2150, Microchem), with a thickness of 350 μm. The wafer was exposed to UV light through the mask and developed in PGMEA (Sigma-Aldrich) in order to produce a mold. PDMS (Sylgard 184, Dow Corning) was subsequently casted on the mold and cured at 80°C overnight. After cutting out the chips, we punched 1 mm inlet and outlet ports. We finally punched a 3 mm hole right downstream of the inlet port. This hole, after being covered with a PDMS piece, acts as a bubble trap. To fabricate channels for the twitching experiments, we followed a similar procedure, but we used a different photoresist (SU8 2025 Microchem) and we adjusted the dimensions of the channel to be 500 μm wide and 100 μm high.

The final assembled chips were obtained by placing the PDMS chips on top of the hydrogel-coated coverslips right after polymerization. This results in a reversible, but sufficiently strong bond between the hydrogel and the PDMS, allowing us to use the chips under flow without leakage for several days. The channels of the chips were filled with M9 medium to keep the hydrogel hydrated for at least 12 hours before being used.

### BACTERIAL STRAINS

Strains used in this work are listed in **Table S2**. All strains were grown in LB medium at 37°C. Overnight bacterial cultures were diluted 1:1000 in fresh LB and grown until mid-exponential phase (optical density at 600 nm: 0.3 to 0.6).

**Table S2.** List of bacterial strains.

| Name | Description | Origin/reference |
| --- | --- | --- |
| PAO1 |  | <sup>10</sup> |
| PAO1 $\Delta flic$ | in-frame deletion of PA1092 | <sup>11</sup> |
| PAO1 $\Delta flic \Delta pilA$ | in-frame deletion of PA1092 and PA4525 | <sup>11</sup> |
| PAO1 $\Delta flic \Delta pilTU$ | in-frame deletion of PA1092, PA0395 and PA0396 | <sup>11</sup> |
| PAO1 mNeonGreen | PAO1 constitutively expressing mNeonGreen (attTn7::miniTn7T-Gm-ptet::mNeonGreen) |  |
| PAO1 mScarlet | PAO1 constitutively expressing mScarlet (attTn7::miniTn7T-Gm-ptet::mScarlet) |  |
| PAO1 GFP | PAO1 constitutively expressing GFP (attTn7::miniTn7T2.1-Gm-GW::PA1/04/03::GFP) |  |

### SINGLE CELL TWITCHING AND ADHESION

Overnight bacterial cultures of PAO1 *Δflic* and PAO1 *Δflic ΔpilTU* were diluted 1:1000 in fresh LB and grown until mid-exponential phase (OD 0.4–0.6). Bacterial cultures were diluted to reach an optical density of 0.4 for PAO1 *Δflic*, and 0.1 for PAO1 *Δflic ΔpilTU*. We then loaded 10 μL of the bacterial culture in the small microfluidic chips (500 μm wide and 100 μm deep) assembled with either hydrogel-coated coverslips or with glass coverslips. We let the bacteria adhere for 30 minutes. We connected the inlet port to a disposable syringe (BD Plastipak) filled with the medium and mounted onto a syringe pump (KD Scientific), using a 1.09 mm outer diameter polyethylene tube (Instech) and a 27G needle (Instech). For twitching experiments, we used a flow of 60 μL·h^−1^ for 5 minutes in order to remove bacteria that did not adhere to the hydrogel surface. We then switched to a flow of 30 μL·h^−1^ before starting image acquisition. Images were taken every 30 s for 1 h.

For adhesion experiments 200 μL of bacterial cultures of PAO1 *Δflic ΔpilA* at OD 0.5 were centrifuged at 6000 rpm and resuspended in 50 μL of LB. We load 10 μL of the bacterial suspension in small microfluidic chips that were hold together with a custom-built clamp to avoid delamination of the chip from the gel under large flow rate. We connected the inlet port as described above and we let the cells adhere for 30 min. We used a flow of 60 μL·h^−1^ for 5 min in order to remove bacteria that did not adhere to the hydrogel surface. Images were then taken every 10 s for 6 min in the center of the channel with a flow of 60 μL·h^−1^ for the first 2 min, 600 μL·h^−1^ for the next 2 and 6000 μL·h^−1^ for the final 2 min.

We aquired images on a Nikon TiE widefield microscope equipped with a Hamamatsu ORCA Flash 4 camera. Images were acquired in phase contrast with a 20× Plan APO NA 0.75 objective.

### BIOFILM FORMATION

Overnight bacterial cultures of PAO1 were diluted 1:1000 in fresh LB and grown until mid-exponential phase. Bacterial cultures were diluted to reach an optical density of 0.05. We then loaded 6.5 μL of the diluted bacterial culture in the big channels (2 mm wide and 350 μm high), from the outlet port. It is important that injected cells do not reach the well of the bubble trap. We let the cells adhere for 30 min before starting the flow. The biofilms were grown at 25°C at a flow rate of 10 μL·min^−1^. For timelapse visualizations of early-stage biofilm formation, we acquired images every 15 min for 15 h in phase contrast with a 40× Plan APO NA 0.9 objective. For the visualization of biofilms, we used a Nikon Eclipse Ti2-E inverted microscope coupled with a Yokogawa CSU W2 confocal spinning disk unit and equipped with a Prime 95B sCMOS camera (Photometrics). We used a 20x water immersion objective with N.A. of 0.95 and z-stacks of the biofilms were taken every 2 μm.

### IMAGE PROCESSING AND DATA ANALYSIS

Snapshot images and movies were generated with Fiji. Images were processed with macros in Fiji and data were analyzed in Python 3 and OriginPro.

#### Single cell twitching

When necessary, drift was corrected using the Correct 3D Drift plugin. Cells were segmented and then tracked using a Trackmate script that tracks spots based on a result table that contains the position of each cell’s center of mass in the movie. Subsequent analysis of cell trajectories was done with a custom Python script. Only tracks with a duration above 10 min were considered. Cell speed was calculated as the net displacement (distance between the last and the first spot of the track) divided by the track duration. An average speed was calculated for each replicate based on at least 50 tracks and displayed values are the average among three replicates and the corresponding standard deviation.

#### Single cell adhesion

Only a portion of the stack with a width equal to half of the channel width taken in the center was considered. Cells were segmented and tracked as above. Only cells attached to the surface from time 0 were considered. We defined the shear stress at the wall of the channel as *σ_s_* = 6*Qμ*/*wh*^2^, where *Q* is the volumetric flow, *μ the viscosity of the fluid and w and h the width and the height of the channel respectively*. Cells attached at time 0 are defined as the number of cells the stays attached for at least 90 s under a shear stress of 0.02 Pa. The percentage of cells that stays attached to the surface is defined as the number of cells attached to the surface just before the next increase of shear stress normalized by the number of cells attached at time 0.

#### Biofilm morphology

Biofilms were imaged after 40 h of growth in proximity to the channel inlet. For each condition we imaged 6 chips (2 chips for each biological replicate) and 30 single colonies were selected and segmented. The radius of the colony is defined as the radius of a circle with an area equal to the substrate area covered by the biofilm. The height of the colony is defined as the maximum height of the orthogonal projection at the center of the colony (found by fitting a circle to the area covered by the biofilm). The displayed height to radius ratio corresponds to all measured colonies.

#### Biofilm spatial organization

For mixing experiments, PAO1 mNeonGreen and PAO1 mScarlet were mixed at a 1:1 ratio before inoculation in the channels. Biofilms were imaged after 40 h of growth at the beginning of the channel. For each condition we imaged 6 chips (2 chips for each biological replicate) and for each chip we acquired around 10 stacks. To correct for the slide tilting we performed a maximum intensity projection of the first 3 slices for each stack. Images were then segmented for the 2 fluorescent channels. To quantify the 1^st^ nearest neighbor distance (1^st^ NND) we selected 100 random pixels in the segmented mNeonGreen picture for each stack. We then calculated the distance between the selected pixels and their nearest neighbor pixels in the corresponding segmented mScarlet picture. This way we obtained an average 1^st^ NND for each stack. For each chip the highest 1^st^ NND value was filtered out and a mean value of the remaining stacks was calculated. The displayed 1^st^ NND corresponds to the average 1^st^ NND for each chip.

#### Antibiotic treatment of biofilms

Biofilms of PAO1 GFP were imaged after 46 h (time 0) of growth at the beginning of the channel. For each condition we imaged 4-5 chips (distributed among 3 biological replicates) and for each chip we acquired 6 stacks. We then switched the medium from LB to LB containing the antibiotic colistin (5 μg/mL, Acros organics) and the dye Propidium Iodide (5 μM, Cayman chemical) for staining of dead cells. Biofilms were imaged every hour in the same positions set at time 0. Stacks acquired at time 0 and after antibiotic treatment were concatenated and the drift was corrected.

To quantify the effect of the antibiotic on the biofilms we measured the volume of live biomass (expressing GFP). Stacks were segmented and the biofilm volume at different times was quantified with the plugin 3D Objects counter and normalized by the value at time 0. We then obtained and average value for each chip. Biofilm volume and surface at time 0 were quantified with the plugin 3D Objects counter. The substrate area covered by the biofilm was measured after performing a maximum intensity projection of the segmented stack. The exposed surface was calculated as the difference between the total surface and the substrate area covered by the biofilm. Exposed surface to volume ratio was calculated for each stack and we obtained an average value for each chip.

To quantify cell death induced by colistin, we selected around 30 - 40 single colonies from stacks acquired after 1 h of antibiotic treatment. Images in the green channel were segmented and used to define the core (C) and the rim (*R*) of the colony by fitting a circle to the area covered by the biofilm. We then performed a radial reslice over 360 degrees in the red channel (dead biofilm) by rotating a line with a length equal to *R* around one of its ends placed in C. We performed an average intensity projection of the resulting stack and measured the intensity profile along a line of length equal to *R* drawn slightly above the plane of contact between the biofilm and the substrate. The intensity of the curves was normalized by the highest value, while the distance was normalized by *R*. We then averaged the curves and calculated the standard deviation for each condition. We then performed integration of the curve between 0 and 0.2 (C) and between 0.8 and 1 (*R*) and we normalized the results by the total area under the curve.

